# Selective and controlled myelin formation by individual interfascicular oligodendrocytes in the mouse corpus callosum

**DOI:** 10.1101/2020.02.20.957522

**Authors:** Tatsuhide Tanaka, Nobuhiko Ohno, Yasuyuki Osanai, Sei Saitoh, Truc Quynh Thai, Kazuya Nishimura, Takeaki Shinjo, Shoko Takemura, Kouko Tatsumi, Akio Wanaka

## Abstract

Single oligodendrocytes produce myelin sheath around multiple axons in the central nervous system. Interfascicular oligodendrocytes (IOs) in the white matter are aligned in rows and facilitate nerve conduction, but their detailed morphologies remain largely unknown. In the present study, we three-dimensionally reconstructed seven IOs in a row in the murine corpus callosum using serial block face-scanning electron microscopy (SBF-SEM). These morphologically polarized IOs extended a thick process with numerous branches from the cytoplasm-rich part of the cell and formed myelin sheaths preferentially around distant axons. The multiple branched processes of each IO myelinated multiple axons having similar diameters with restricted myelin thicknesses, indicating that individual IOs have their own myelination profiles even on distinct target neurons. Consistent with the finding, the IOs transduced and visualized with the rabies viral vector expressing GFP showed statistically significant variations in the myelination patterns. We further reconstructed the sheath immediately adjacent to those derived from the seven IOs; the thicknesses of both sheaths were significantly correlated despite emanating from different IOs. These results proposed a rule that myelination by individual IOs, regulated by interaction with ensheathed axons, is selective in specific axons and at the same time orchestrated at the whole cell level.

**Main points:** Callosal oligodendrocytes form myelin sheaths preferentially around distant axons.

Each oligodendrocyte myelinate multiple axons having similar diameters with restricted myelin thicknesses.

The thicknesses of adjacent myelin sheaths are similar.

## Introduction

Myelin facilitates the rapid propagation of electrical impulses along axons by saltatory conduction. Oligodendrocytes extend multiple processes and form a myelin sheath around axons in both the gray and white matter of the central nervous system (CNS) (Kahn et al., 1986). The formation of myelin can modulate complex neuronal interconnection, and therefore abnormal myelination often results in neurological disorders and mental illnesses (Baumann and Pham-Dinh, 2001), (Hildebrand et al., 1993), (McKenzie et al., 2014), (Liu et al., 2012), (Wake et al., 2011). Recent studies using selective fluorescence labeling and three-dimensional (3D) reconstruction of electron microscopic images suggest that some oligodendrocytes preferentially ensheath excitatory or inhibitory axons (Osanai et al., 2017), (Zonouzi et al., 2019). Given that oligodendrocyte depolarization at the whole-cell level can modulate nerve conduction by ensheathed axons (Yamazaki et al., 2014), oligodendrocytes may dynamically regulate nerve conductivity of specific neural circuitries through selective myelin formation. However, the mechanisms of selection as well as the patterns of regulation are still largely unknown.

While oligodendrocytes are widely regarded as being functionally homogeneous throughout the CNS, they are morphologically heterogeneous (Mori and Leblond, 1970). In addition, recent analyses of transcriptional profiles showed remarkable heterogeneity of oligodendrocytes in adult murine brain (Marques et al., 2016). Oligodendrocytes in the white matter of the CNS are often aligned in rows between the nerve fibers, and are called interfascicular oligodendrocytes (IOs) (Kahn et al., 1986), (Suzuki and Raisman, 1992), (Ogawa et al., 2001). Although these linearly arranged IOs may collectively regulate nerve conduction of specific groups of neurons, the heterogeneity of the grouped IOs in terms of axonal selection and myelin sheath formation remains to be elucidated.

In this study, using 3D reconstructions of serial electron microscopic images in a large volume of corpus callosum in adult mice, we traced the cell soma, processes and myelin sheaths of a row of seven IOs. All seven reconstructed IOs extended their processes and formed myelin sheaths around distant axons. Each IO had its own myelination pattern represented by myelin thicknesses and the diameters of ensheathed axons. These myelination patterns, unique to individual IOs, showed significant variation among IOs, and this finding was further confirmed by morphometric analysis of myelinated axons using rabies virus-mediated oligodendrocyte labeling. We further reconstructed the myelin sheath (presumed to be formed by an IO other than any of the seven targeted IOs) adjacent to the originally reconstructed sheath and found that their thicknesses are significantly correlated. Taken together, our data indicate that oligodendrocytes in the corpus callosum exhibit diverse properties in terms of their structural interactions with axons. This contradicts the common belief that oligodendrocytes constitute a non-diverse population.

## Materials and Methods

### Animals

All animal protocols were approved by the Animal Care Committee of Nara Medical University in accordance with the policies established in the NIH Guide for the Care and Use of Laboratory Animals. Female 8-week-old C57BL/6J mice were used in all experiments.

### Immunohistochemistry

Mice were anesthetized and perfused transcardially with saline followed by 4% paraformaldehyde (PFA) in 0.1 M phosphate buffer (pH 7.4) (PB). Brains were removed, postfixed overnight in the same fixative, and then immersed in 30% sucrose in PB overnight. Next, the brains were frozen in powdered dry ice, embedded in Tissue-Tek OCT compound (Sakura Finetek, Torrance, CA, USA), and stored at −80°C prior to sectioning. Twenty-micrometer-thick transverse sections were cut on a cryostat and mounted onto glass slides. Sections were immersed in PB containing 5% bovine serum albumin and 0.3% Triton X-100 for 1 h. Antibodies against adenomatous polyposis coli (1:250; APC or CC1, Merck Millipore, Billerica, MA, USA) or GFP (1:8000, Nacalai Tesque, Kyoto, Japan) were applied overnight at 4°C. Alexa Fluor 488-conjugated IgG (Life Technologies, Grand Island, NY, USA) was used as secondary antibody. Images were captured using a confocal laser scanning microscope with Fluoview FV1000-D software (Olympus, Tokyo, Japan).

### Serial block face scanning electron microscopy (SBF-SEM)

Mice were anesthetized and perfused transcardially with saline followed by 4% PFA and 2.5% glutaraldehyde in PB. Brains were removed and postfixed overnight in the same fixative. The brains were cut into 300-μm-thick sagittal slices with a vibratome slicer. Slices were then treated with 2% OsO_4_ (Nisshin EM, Tokyo, Japan) in 0.1 M cacodylate buffer (CB) containing 0.15% K_4_[Fe(CN)_6_] (Nacalai Tesque), washed four times with CB, and incubated with 0.1% thiocarbohydrazide (Sigma Aldrich) for 20 min and 2% OsO_4_ for 30 min at room temperature (RT). The slices were then treated with 2% uranyl acetate at 4°C for 12 h and at RT for 30 min, and stained with Walton’s lead aspartate at 60°C for 30 min. The slices were dehydrated by passing the tissue through an ethanol serie (60% – 80% – 90% – 95% – 100%) at 4°C, infiltrated sequentially with acetone dehydrated with a molecular sieve, a 1:1 mixture of resin and acetone, and 100% resin, and then embedded with Aclar film (Nisshin EM, Tokyo, Japan) in Epon 812 epoxy resin with carbon (Ketjen black). The specimen-embedding resin was polymerized at 40°C for 6 h, 50°C for 12 h, 60°C for 24 h and finally at 70°C for 2 days. After trimming of the region containing the corpus callosum from the brain, the samples were imaged with a Merlin (see below) equipped with the 3View system and an OnPoint backscattered electron detector (Gatan, Pleasanton, CA, USA).

Merlin (Carl Zeiss, Munich, Germany) is a field emission-type scanning electron microscope with a single electron beam. The beam was set to 1.5kV acceleration voltage, 130 pA beam current and 6.7 mm working distance to ensure high-quality image acquisition. The imaging conditions of the OnPoint backscattered electron detector (Gatan) were 5.7 nm/pixel and 0.5 μs/pixel dwell time. The single-slice size was X-Y = 17,408 pixels x 17,408 pixels and Z thickness = 40 nm. Three-dimensional cubic images were reconstructed from 2,000 slices (see below), corresponding to the field as a field of X-Y = 99.2256 μm x 99.2256 μm and Z = 80 μm. The estimated electron dose was 12.48 e-/nm^2^.

### Observation of SBF-SEM images

Serial images of SBF-SEM were handled with Fiji/ImageJ and segmented using Microscopy Image Browser (http://mib.helsinki.fi/) and reconstructed to 3D images with Amira software (Thermo Fisher Scientific). Axonal diameters were calculated from length and surface area data, assuming axons to be cylinders (Ohno et al., 2011). Axonal length was calculated using Amira software as the path of the center point of the minimum inscribed sphere of an axonal profile.

### Injection of viral vectors into the cortex and corpus callosum

Viral vectors and injection procedures were described in detail previously(Osanai et al., 2017), (Osanai et al., 2018). Briefly, mice were anesthetized with an intraperitoneal injection of ketamine/xylazine solution (125 mg/kg and 10 mg/kg, respectively) and placed in a stereotaxic frame (Narishige, Tokyo, Japan). AAV-DsRed2 or AAV-BFP was stereotaxically injected into either the motor cortex (1.0 mm posterior and 0.8 mm lateral to the bregma, at a depth of 0.3 mm) or the somatosensory cortex (1.0 mm posterior and 1.5 mm lateral to the bregma, at a depth of 0.3 mm). After opening the skull over the injection site, 1 μL of each AAV vector solution (1.0 × 10^9^ viral genomes) was injected through pulled glass pipettes (inner diameter 20–30 μm) using an air pressure system, which took about 3 min. Two weeks after the initial injection, 1 μL of RV-GFP (5.3 × 10^2^ IU) was stereotaxically injected into the corpus callosum (1.0 mm posterior and 0.5– 0.8 mm lateral to the bregma, at a depth of 1.0 mm). The mice were perfused with saline followed by 4% PFA in PB 4 days after RV-GFP injection. The brains were dissected out and subjected to confocal imaging (Osanai et al., 2017), (Osanai et al., 2018).

### Statistical analysis

Data are presented as mean ± SEM. Statistical analyses were performed using one-way analysis of variance for multiple groups. Pearson correlation coefficienr test was performed for two groups. All *n* is indicated in the figure legends.

## Results

### Beads-like cell cluster of IOs with asymmetric process extension

IOs are aligned in rows between the nerve fibers in the white matter of the CNS (Kahn et al., 1986), (Suzuki and Raisman, 1992), (Ogawa et al., 2001). Indeed, in the corpus callosum of adult mice, cells displaying nuclear DAPI staining and immunostaining for CC1, a marker of mature oligodendrocytes, formed chains that we refer to as beads-like cell clusters (Fig. 1A, B). CC1-negative cells in these images are likely to include astrocytes, microglia, and endothelial cells, which are also found in the white matter (Sampaio-Baptista and Johansen-Berg, 2017). In order to follow the beads-like cell clusters in the corpus callosum at higher resolution and in three dimensions, we employed serial block face scanning electron microscopy (SBF-SEM). We acquired 2000 serial images (40 nm thickness/image) spanning a total volume of 95.7 × 96.7 × 80 μm3, at a resolution of 5.7 nm/pixel (Fig. 1C, Supplementary movie 1). In this volume, we first traced the nucleus of all cells. Oligodendrocytes, characterized by their nuclear morphology, were aligned in a row among bundles of myelinated axons (Peters et al., 1991) (Fig. 1D, E). We focused seven oligodendrocytes (O1-O7) and traced the cytoplasm of oligodendrocytes which formed a beads-like cell cluster in the corpus callosum (Fig. 1E, F, Supplementary movie 2).

**Fig.1.**
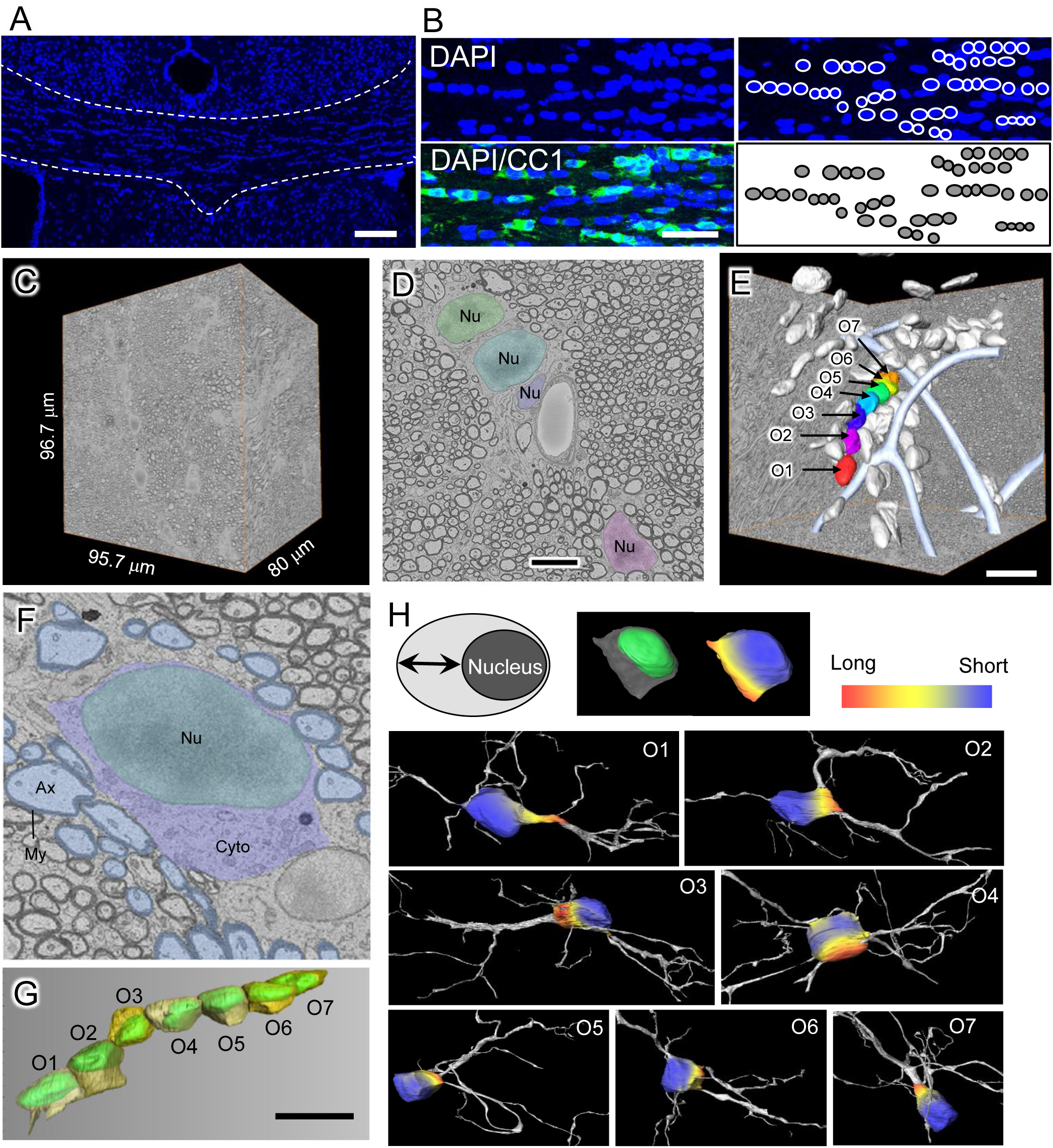
Beads-like cell clusters of interfascicular oligodendrocytes. **A**: Confocal image of the corpus callosum stained with DAPI (blue). Dashed lines demarcate the corpus callosum. Scale bar, 100 μm. **B**: Left panels, Confocal images of the corpus callosum stained with DAPI (blue) and CC1 (green). Scale bar, 50 μm. Right panels, beads-like cell clusters in the left panel are marked by white circles (upper) and each cluster is colored (lower). **C**: Observation area of serial block face scanning electron microscopy (SBF-SEM). The data set spans a 95.7 × 96.7 × 80 μm^3^volume and was sectioned at 40 nm thickness. **D**: A representative EM image of the corpus callosum. Note that the nuclei (Nu) of oligodendrocytes make a beads-like cluster in this field. Scale bar, 10 μm. **E**: Three-dimensional renderings of SBF-SEM images of seven oligodendrocytic (O1 to O7) nuclei, which we focused on in the present study. Tubular structure (light blue) indicate blood vessel. Scale bar, 20 μm. **F**: A representative EM image of an oligodendrocyte in the corpus callosum. The nucleus (Nu) is biased to one side of the cell and the cytoplasm (Cyto) is predominantly on the opposite side. Ax, axon (light blue); My, myelin. **G**: Three-dimensional reconstructed image of the O1 to O7 oligodendrocytes (indicated in **E**), highlighting the shapes and locations of nuclei (green) and cytoplasm (yellow). Note that the nuclei are all biased to one side of the cells, as seen in **F**. Scale bar, 10 μm. **H**: Distance between the nuclear and plasma membranes is color-coded (blue, short; red, long). The upper left panel shows a diagram of an oligodendrocyte. The upper center panel shows representative 3D images of the cell body of an oligodendrocyte, highlighting the shape of the nucleus (left, green) and the inter-membrane distance (right, color-coded). Lower, three-dimensional rendering of the seven oligodendrocytes (O1 to O7) with color coding. Note that thick processes protrude from the cytoplasm-rich area (red). Processes of each oligodendrocyte are in gray.

Oligodendrocytes have been considered as having a narrow cytoplas (Cervos-Navarro et al., 1987). However, we found that all the reconstructed oligodendrocytes had partially wide cytoplasm containing various organelles, and the position of the nucleus was biased in different directions (Fig. 1F, G). To investigate the relationship between cell polarity and process formation, we measured the distance between the nuclear and plasma membranes, and reconstructed all processes extending from the cell bodies. In each IO, a single thick process with multiple branches protruded from regions of the cell where the cytoplasm was widest (Fig. 1H). On the other hand, only a few very thin processes extended from sites with narrow cytoplasm (Fig. 1H). These results suggest that oligodendrocytes possess a previously unappreciated polarity associated with asymmetric extension of thick processes from the cell bodies.

### Distinct myelin profiles are formed by individual IOs in the corpus callosum

Individual oligodendrocytes form multiple myelin sheaths at the tips of the processes extended from their cell bodies (Simons and Nave, 2016). We next followed these processes and identified the myelin sheaths formed by the reconstructed oligodendrocytes. Identification of the myelin sheaths formed by individual processes of particular oligodendrocytes was difficult because processes frequently made contact with compact myelin sheaths that just passed nearby and were not necessarily formed by those processes. To overcome this difficulty, we focused on the presence of an outer tongue, a cytoplasmic domain that runs continuously along the myelin sheath and is directly connected to the oligodendrocyte process (Chang et al., 2016) (Fig. 2A, B). By confirming the connection of individual processes to such outer tongues (Fig. 2B, arrowheads), we identified the specific compact myelin sheaths that were formed by processes originating from the reconstructed cell bodies of individual IOs (Fig. S1, arrowheads). We identified a total of 91 myelin sheaths formed by the seven IOs, each of which had many branched processes (Fig. 2C, Supplementary movie 3–10). All the IOs extended long processes to distant axons even though there were many axons in closer proximity (Fig. 2C and Fig. 1F). The average linear distances between cell bodies and myelin sheaths were all within the range of 10 – 20 μm, with no significant difference among the IOs (Fig. 2D; *p* = 0.9, one-way ANOVA) and no obvious relationship between the linear distance and the IO cell body volume (Fig. 2E). Tracing the processes to the outer tongues enabled reliable identification of distant myelin sheaths formed by individual oligodendrocyte processes in the white matter.

**Fig. 2.**
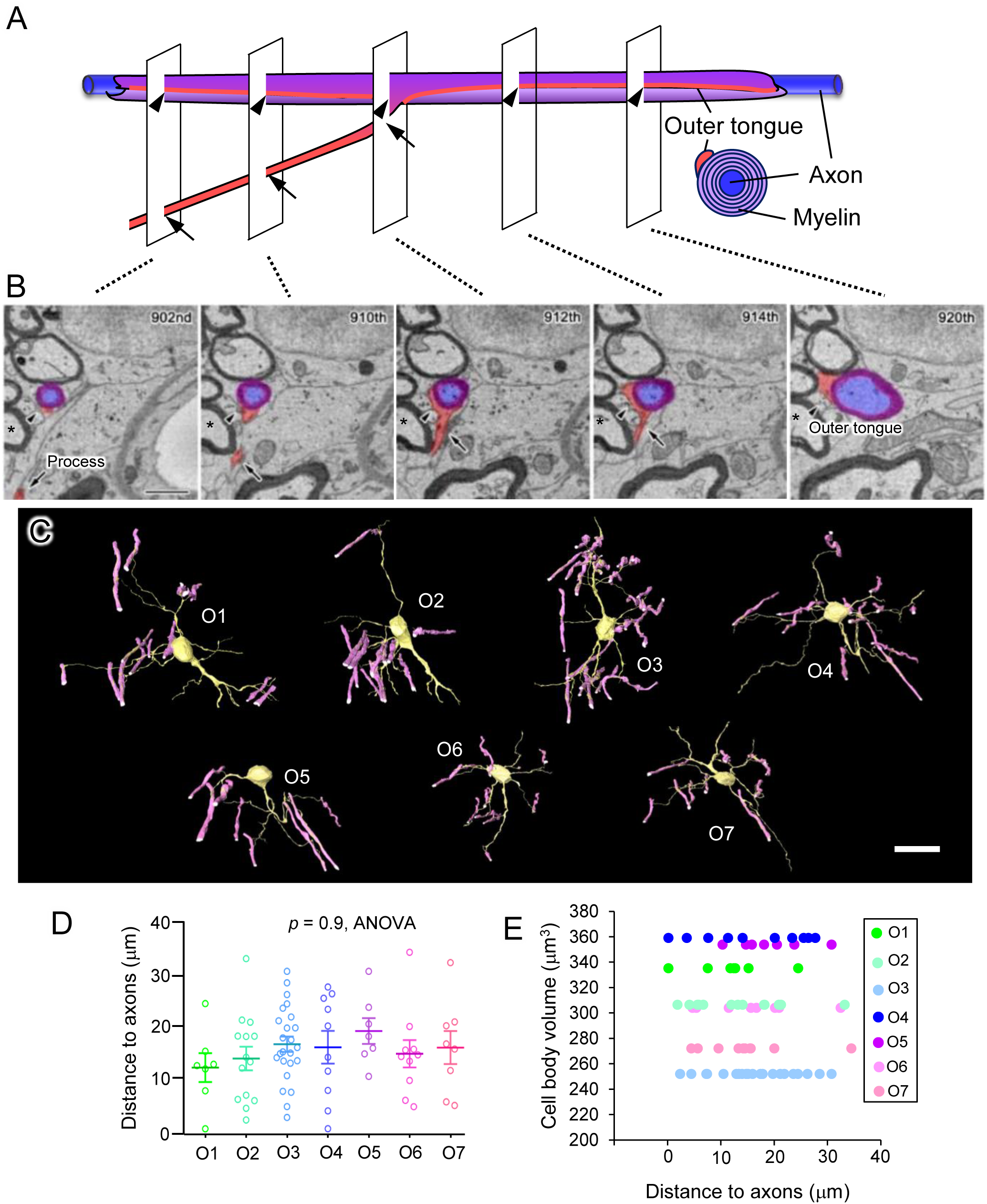
Three-dimensional structural characteristics of oligodendrocytes in the corpus callosum. **A**: Diagrams show a 3D image and a cross section of an axon and a myelinating oligodendrocytic process. Five planes correspond to the EM images shown in **B**. An oligodendrocyte process (orange, arrows) approaches an axon (blue) ensheathed by myelin (purple) and attaches. A cytoplasmic domain, the outer tongue (orange, arrowheads), runs along the myelin sheath. **B**: Serial EM images of an oligodendrocytic process and its targeted axon. Arrows, process of oligodendrocyte; arrowheads, outer tongue (orange). Blue, targeted axon. Purple, myelin. Scale bar, 1 μm. Note that the process passes by a myelinated axon (asterisk) but does not form myelin with an apposed outer tongue. **C**: Three-dimensional rendering of seven oligodendrocytes (O1 to O7). Yellow, cell body and processes; magenta, myelin sheath. Scale bar, 20 μm. **D**: Quantification and statistical analysis of the distances between cell bodies and myelin sheaths formed by the seven oligodendrocytes in the beads-like cell cluster. One-way ANOVA, *p* = 0.9. Data are presented the mean ± SEM. **E**: Correlation between the cell body volume and the lengths of processes for the seven oligodendrocytes. There is no positive correlation. D-E, n = 7 (O1), 14 (O2), 24 (O3), 10 (O4), 7 (O5), 10 (O6), 8 (O7).

The size of unmyelinated axons varies along their trajectories, partly due to the presence of specialized structures such as varicosities. In the serial images of a myelinated nerve fiber (Fig. 3A), we noticed that the axon diameters also significantly varied, sometimes up to several-fold, along the entire length examined (Fig. 3A–C). This finding implied that relying on an axonal diameter obtained from a single section could cause unwanted bias or variability. Ideally, one would obtain the average axonal diameter from cross sections of the axons in all the traced images, but we could not always see cross sections, depending on the direction of axonal projection. As an alternative approach, we measured the surface area and length of an axon with Amira software and calculated the average axonal diameter (Ohno et al., 2011) (Fig. 3B, C). In the axon, whose cross section was obtained over its entire length in the observed area, the average axonal diameter calculated from the perimeter of each axonal cross section was almost identical to the mean axonal diameter calculated from the surface area of the axon and quite distinct from the median axonal diameter (Fig. 3B, C).

**Fig. 3.**
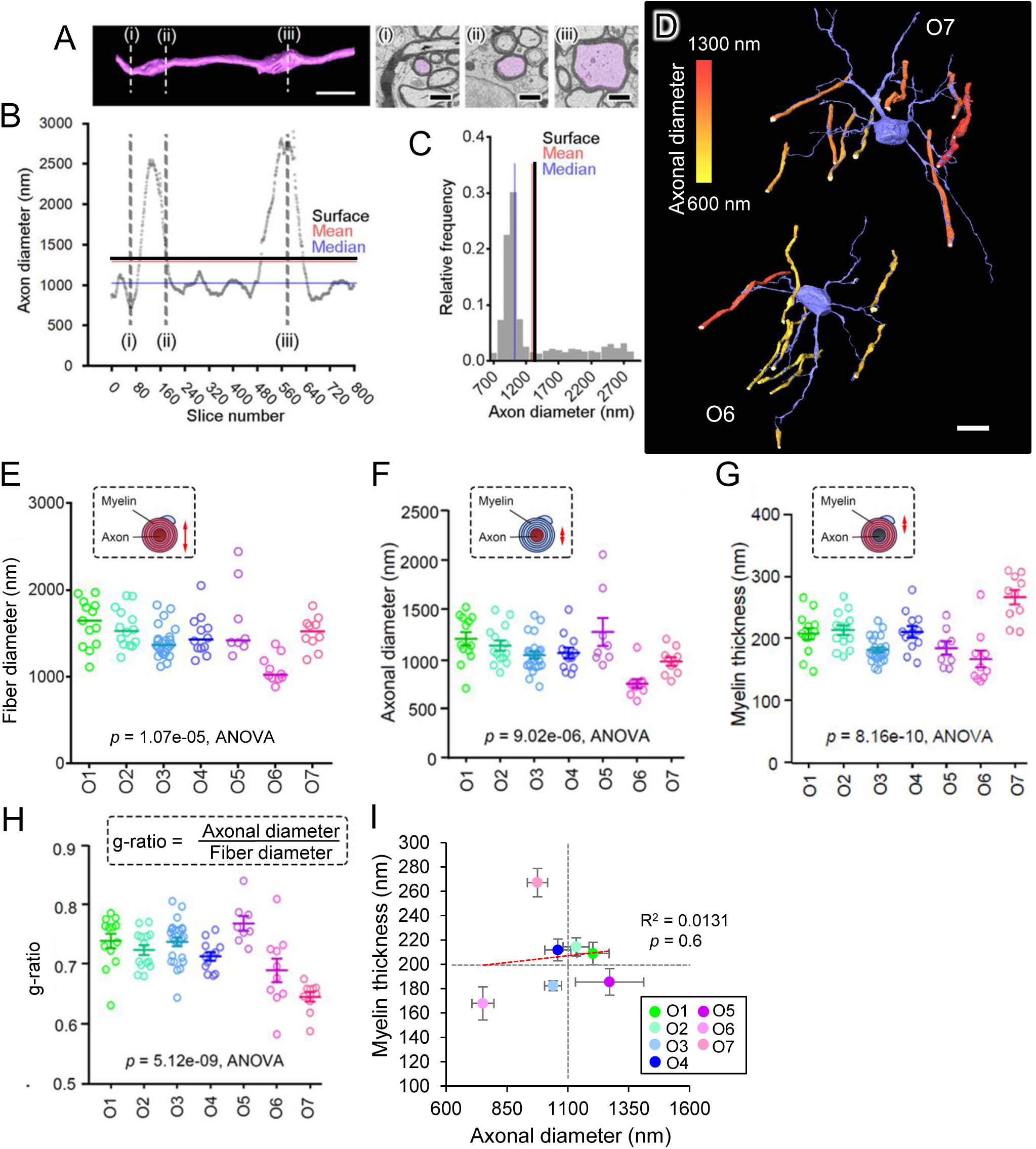
Distinct myelin profiles of individual oligodendrocytes in beads-like cell clusters. **A**: A representative three-dimensional rendering of an axon (magenta) is shown in the left panel and (i), (ii) and (iii) indicate the regions shown in the EM cross sections (right panels). Scale bar, 5 μm (left panel), 1 μm (right panel). **B**: The diameters of the axon were plotted over its entire length (covered by 800 slices). Dashed lines correspond to (i), (ii) and (iii) in **A**. Red and blue lines indicate the mean and median values, respectively, of axonal diameters obtained from all the obtained images. The black line (Surface) indicates the mean axonal diameter calculated from the surface area of the axon. **C**: Histogram of the relative frequencies of the axon diameters indicated in **B**. Red, blue and black lines are the same as those in **B**. Note that the mean axonal diameter calculated from the surface area of the axon was much more similar to the mean axon diameter than to the median axon diameter. **D**: Mean diameters of the axons targeted by two representative oligodendrocytes (O6 and O7) are color-coded (yellow, is thin; red, thick). Blue, cell bodies and processes of the oligodendrocytes. Scale bar, 10 μm. **E**: Quantification of the fiber diameters formed by each oligodendrocyte. Variation among the seven oligodendrocytes was statistically significant (*p* = 1.07e-5, one-way ANOVA). **F**: Quantification of the diameters of axons myelinated by each oligodendrocyte. Variation among seven oligodendrocytes was statistically significant (*p* = 9.02e-06, one-way ANOVA). Data are presented the mean ± SEM. **G**: Myelin thicknesses of the sheaths formed by seven oligodendrocytes varied significantly (*p* = 8.16e-10, one-way ANOVA). Data are presented the mean ± SEM. **H**: The g-ratios of the fibers myelinated by the seven oligodendrocytes showed statistically significant variation (*p* = 5.12e-09, one-way ANOVA). Data are presented the mean ± SEM. **I**: Relationships between axonal diameters (x axis) and myelin thicknesses (y axis) for each oligodendrocyte. Note that the oligodendrocytes have their own patterns of myelination, and these patterns do not simply follow the rule “a larger axon has a thicker myelin sheath”. Data are presented the mean ± SEM. The red dashed line represents the regression line. None of the correlations were statistically significant. R^2^; Decision coefficirnt. E-I, n = 13 (O1), 14 (O2), 24 (O3), 12 (O4), 8 (O5), 10 (O6), 10 (O7).

Reliable identification of the myelin sheaths formed by individual oligodendrocyte processes (see above) and estimation of mean fiber diameters allowed us to analyze structural properties of myelin sheaths produced by individual IOs and their ensheathed axons. Interestingly, the measurements of fiber diameters (myelin + axon) revealed that the fiber diameters of axons ensheathed by each IO were restricted to a certain range (Fig. 3D). We found that there were differences in fiber diameters, axon diameters, myelin thicknesses and g-ratios (fiber diameter/axon diameter) of myelin sheaths derived from individual IOs (Fig. 3E–H; *p* < 0.001, one-way ANOVA). Myelination patterns can be allotted into four quadrants: thin myelin–small (diameter) axon, thin myelin–large (diameter) axon, thick myelin–small axon, thick myelin–large axon (Fig. 3I). For example, O5 formed thin myelin around a large axon (myelin thickness, 185.5 ± 11.0 nm; axon diameter, 1271.5 ± 140.5 nm; g-ratio, 0.77 ± 0.012), while O6 formed thin myelin around a small axon (myelin thickness, 167.9 ± 13.6 nm; axonal diameter, 750.7 ± 45.1 nm; g-ratio, 0.69 ± 0.02). O7 formed especially thick myelin around a small axon (myelin thickness, 267.1 ± 11.6 nm; axon diameter, 974.9 ± 41.8 nm; g-ratio, 0.65 ± 0.008). Variations in the thickness of myelin, the diameter of axons surrounded by myelin and g-ratios in individual oligodendrocytes were biased to a narrow range (Fig. 3F-I). We observed no correlation between the axonal diameter and myelin thickness (Fig. 3I). These results indicate that IOs comprise a heterogeneous population that is endowed with different patterns of myelination, underlining a clear contrast with the peripheral nervous system where myelin thickness has a linear correlation with axonal diameter (Gillespie and Stein, 1983).

We observed no correlation between the distance from cell bodies to myelinated axons and myelin thickness (Fig. S2A, B). In addition, there was no correlation between the distance from cell bodies to myelinated axons and axonal diameter (Fig. S2C, D). Next, we hypothesized that the cell body volume of the IOs might affect the myelin patterns that we observed. To investigate this, we performed a correlation analysis and found no correlation between the soma size of an oligodendrocyte and either myelin thickness or myelinated axonal diameter (Fig. S2E-H). Although each oligodendrocyte in a beads-like cell cluster forms long processes, these results indicate that neither the length of the processes nor the size of the cell body affect the myelination pattern.

Three-dimentional reconstruction of serial EM images is a very powerful tool to analyze detailed structures of oligodendrocytes, but at the same time there is a technical limitation: we cannot analyze oligodendrocytes in a way that is amenable to statistical analysis due to complexities of cellular mophologies. To compensate for this limitation, we employed genetic labeling with a rabies virus (RV) vector. Injection of the RV harboring a gene encoding GFP (RV-GFP) into the mouse white matter can sparsely label oligodendrocytes (in sufficient numbers for statistics) and can visualize the morphology of individual oligodendrocytes in the white matter (Osanai et al., 2017). We first injected AAV-DsRed2 and AAV-BFP into different hemispheres to label callosal axons connecting the bilateral hemispheres of the cerebral cortex (Fig. 4A–C). We next injected RV-GFP into the corpus callosum and examined the oligodendrocyte processes wrapped around axons labeled with either DsRed2 or BFP, using Z-series confocal images. We measured the diameters of fibers (axon + myelin). As was the case in SBF-SEM, we also found that there were differences in the diameter of myelin formed by individual oligodendrocytes (Fig. 4D; *p* < 0.001, one-way ANOVA). This variation suggests that each oligodendrocyte has its own characteristic myelination pattern and/or its own preferred axon diameter. We also observed no correlation between the distance from cell bodies to myelinated axons and fiber diameter (Fig. S3). These results further support the concept that each oligodendrocyte has its charactetistic myelination pattern and its own preference of axon diameters, which are independent of the sizes of its cell body and processes.

**Fig. 4.**
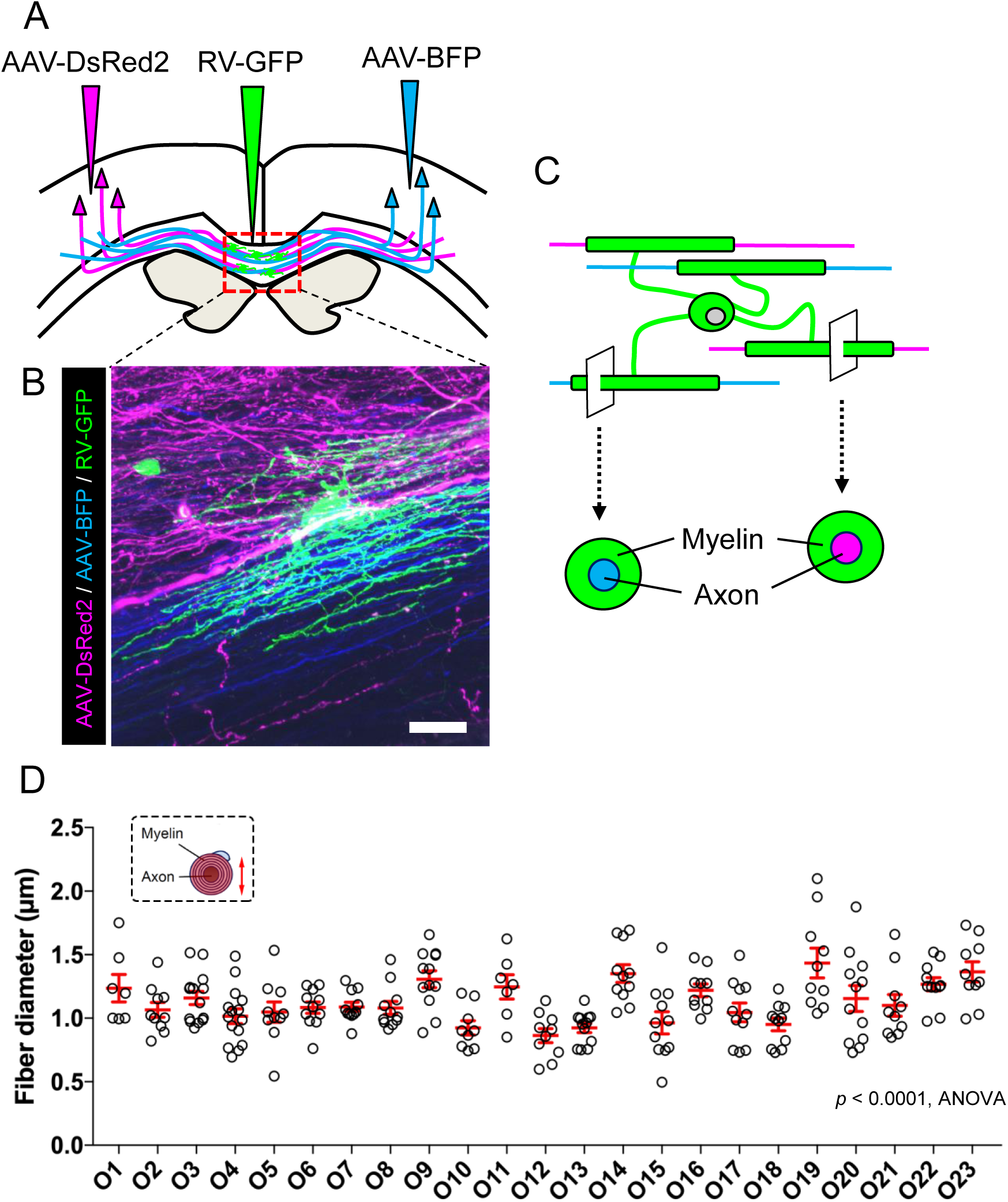
Rabies virus-mediated oligodendrocyte labeling. **A**: Schematic representation of viral injections into each hemisphere and the corpus callosum. **B**: A representative magnified confocal micrograph showing a RV-GFP-positive oligodendrocyte (green) and axons labeled with AAV-BFP (blue) or AAV-DsRed2 (red) in the corpus callosum. Scale bar, 20 μm. **C**: Schematic representation of myelinated axons labeled by AAV-BFP or AAV-DsRed2 and a myelinating oligodendrocyte process (green). **D**: Quantification of the fiber (axon + myelin sheath) diameter formed by individual oligodendrocytes. The diameters of the fibers myelinated by 23 oligodendrocytes (OL1–OL14 are myelinated fibers derived from the somatosensory cortex and OL15–OL23 are myelinated fibers derived from the somatosensory cortex and motor cortex) were analyzed with one-way ANOVA and the variation among fibers was significant (*p* < 0.0001). Data are presented the mean ± SEM. n = 7 (O1), 10 (O2), 14 (O3), 15 (O4), 11 (O5), 11 (O6), 11 (O7), 11 (O8), 12 (O9), 9 (O10), 6 (O11), 10 (O12), 12 (O13), 11 (O14), 11 (O15), 10 (O16), 11 (O17), 11 (O18), 10 (O19), 12 (O20), 10 (O21), 11 (O22), 10 (O23).

### Myelin thickness and axonal diameter are similar in adjacent internodes

Our finding that myelin thickness variable among individual oligodendrocytes ensheathing axons with similar diameter may argue against the notion that myelin thickness is associated with axonal sizes (Hildebrand and Hahn, 1978), (Gillespie and Stein, 1983). In order to address this potential discrepancy, we asked whether adjacent myelin profiles on single axons are similar, given that none of the 91 myelin sheaths we reconstructed from 7 IOs were situated on the same axon. We therefore investigated the myelin profiles of sheaths adjacent to the sheaths formed by O1-O7 oligodendrocytes (Fig. 5A-D). Correlation analysis of axonal sizes revealed a significant correlation between the axonal diameters in two adjacent internodal segments covered by the myelin of reconstructed oligodendrocytes (x axis) and adjacent myelin sheath (y axis) in general (Fig. 5E, R^2^ = 0.586, p<0.001). The correlation of axonal sizes in the adjacent internodal segments is considered significant even when the axonal sizes are classified by each oligodendrocyte (Fig. 5F, median R^2^= 0.674), since the correlation was mostly diminished when the axonal sizes are randomly shaffled (Fig. 5G, median R^2^ = 0.056 in this randomization), and the histogram of median R^2^ following 1000 randomization trials only 2 trials showed the median R^2^ higher than that in the actual dataset (Fig. 5H). These data demonstrate that the sizes of axons are kept consistent in adjacent intermodal segments. Importantly, correlation of the myelin thickness in two adjacent internodal segments covered by the myelin of reconstructed oligodendrocytes (x axis) and adjacent myelin sheath (y axis) was also significant in general (Fig. 5I, R^2^= 0.298, p<0.001). Furthermore, the correlation of myelin thickness in adjacent internodal segments is considered significant when the myelin thickness is classified by each oligodendrocyte (Fig. 5J, median R^2^=0.710), since the correlation was mostly diminished when the myelin thickness were randomly shaffled (Fig. 5K, median R^2^ = 0.033 in this randomization), and the histogram of median R^2^ following 1000 randomization trials only 1 trial showed the median R^2^ higher than that in the actual dataset (Fig. 5L). Collectively, these results demonstrate that the myelin thickness as well as sizes of axons are kept consistent in adjacent internodes and suggest that individual oligodendrocytes selectively ensheath groups of axons which requires specific thickness of myelin sheath.

**Fig. 5.**
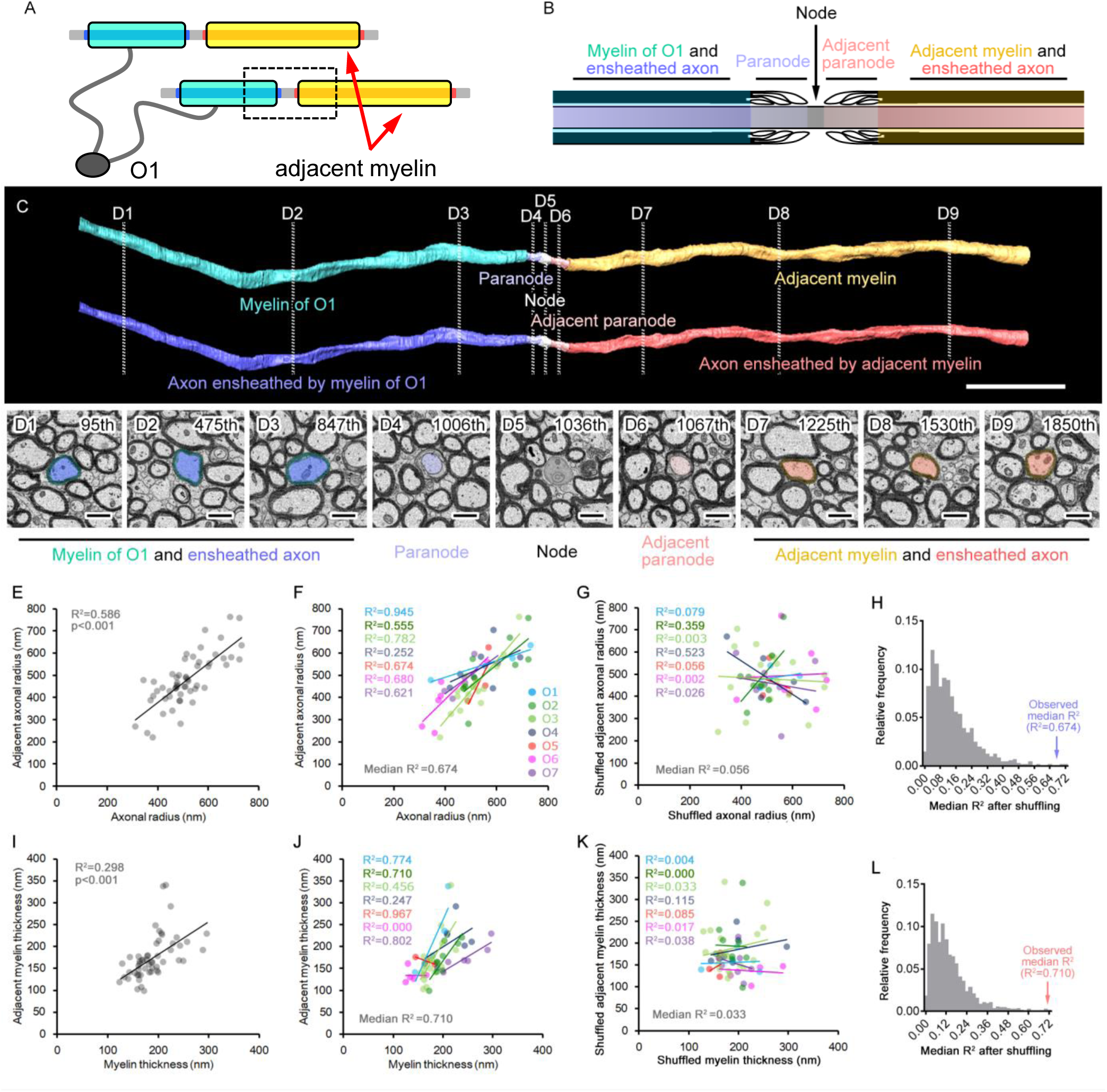
Myelin thickness and axonal diameter are similar in adjacent internodes. **A**: Schematic representation of axons and ensheathing myelins (blue) from a 3D-reconstructed oligodendrocyte (O1) with adjacent myelin sheaths (yellow) from a different oligodendrocyte. **B**: Node of Ranvier (gray) with a flanking paranode (light blue) and an adjacent paranode (pink), located between the myelin of O1 (blue green) with its ensheathed axonal segment (blue) and the adjacent myelin (orange) with its ensheathed axonal segment (red). **C, D**: Three-dimensional reconstruction of an axon myelinated by O1 (**C**; colors are the same as in **A** and **B**), and transverse electron microscopic images (**D**1-9) corresponding to the positions indicated in **C** (dashed lines). Myelin (**C**, upper image) and the ensheathed axon (**C**, lower image) are shown separately. Images **D**1-3 were retrieved from the internodal segment of O1; images **D**4-6 correspond to the paranode, node and adjacent paranode, respectively; and images **D**7-9 are derived from the internodal segment of adjacent myelin. **E, F**: Correlation of the axonal radii in the internodal segments ensheathed by the myelin of segmented oligodendrocytes (x axis) and adjacent axonal radii (y axis), in total (**E**) or grouped for each oligodendrocyte (**F**; O1 to O7 are in different colors). **G**: Correlation of the axonal radii after random shuffling. **H**: Frequency distribution of the median R^2^ for correlation of the axonal diameters after 1000 random shufflings and the observed median R^2^(blue arrow). **I, J**: Correlation of the myelin thicknesses of segmented oligodendrocytes (x axis) and adjacent myelin sheaths (y axis), in total (**I**) or classified by each oligodendrocyte (**J**; colors are the same as in **F**). **K**: Correlation of the myelin thicknesses after random shuffling. **L**: Frequency distribution of the median R^2^ for correlation of the myelin thickness after 1000 random shufflings and the observed median R^2^(red arrow). R^2^; Decision coefficirnt. Bars: 10 μm (**C**) or 1 μm (**D**1-9). E, I, n = 54. F, G, J, K, n = 4 (O1), 9 (O2), 20 (O3), 7 (O4), 3 (O5), 5 (O6), 6 (O7).

## Discussion

To date, high-resolution features of neurons and glial cells have been obtained through dye-filling (Nimmerjahn et al., 2004), (Nimmerjahn et al., 2005) or virus-mediated genetic labeling (Ehrengruber et al., 2003), (Haber et al., 2006), but these techniques are inadequate for comprehensive analyses of relatively large volumes. Recent advances in automated imaging as well as data processing have paved a new way for reconstruction of complete neuron–glial cell connections in a large volume of the nervous system, which will provide key insights into the functions of the glial cells (Tomassy et al., 2014), (Ohno et al., 2014), (Morizawa et al., 2017), (Yamasaki et al., 2014), (Duncan et al., 2018), (Snaidero et al., 2014), (Helmstaedter et al., 2013), (Kasthuri et al., 2015), (Briggman KL, 2012). In the present study, we found that axonal diameter changed even within a single fiber, using SBF-SEM of large volumes of tissue (Fig. 3A). Taking this finding into account, g-ratios calculated from a conventional electron microscopic image may not represent the average data for a myelinated axon, but biased values. To circumvent such possible bias, we extrapolated the mean diameter of an axon from the surface area and employed the values for analyses of oligodendrocyte–neuronal interactions (Fig. 3B, C).

Another structural characteristic found in the present study is of the cell soma of IOs. The position of the nucleus in all cell bodies we observed was biased, indicating that oligodendrocytes are highly polarized cells, and thick processes were formed from areas where the cytoplasm was widest. It has been reported that oligodendrocyte cell bodies are not biased (Cervos-Navarro et al., 1987); however, our data contradict this. Myelin sheath formation requires sorting and trafficking of myelin components from the cell body (Maier et al., 2008). Oligodendrocytes show a polarized phenotype in culture, where they form sheet-like membranes in which myelin proteins and lipids are localized (Dubois-Dalcq et al., 1986). This implies that the sorting and trafficking of myelin components from the *trans*-Golgi network to the myelin-like membrane is an intrinsic property of oligodendrocytes under *in vitro* conditions. We found that the nucleus lies in an eccentric position in each oligodendrocyte; a large mass of cytoplasm occurs at one pole of the cell in beads-like cell clusters, and thick processes extended from the polarized large mass of cytoplasm. These results indicate that polarized cells are generated in the beads-like cell cluster. The locations of nuclei differ from cell to cell in the beads-like structure. Therefore, the elongation of thick processes and myelin formation may occur in all directions from the whole beads-like cell cluster. As shown in Fig. 1G, the oligodendrocytes in the cluster are in contact with each other. It has been reported that cell polarity is triggered by cell–cell adhesion (Desai et al., 2009), (Dupin et al., 2009). Oligodendrocytic cell–cell contacts in beads-like cell clusters may be the regulator of nucleus positioning and intracellular polarized organization.

In addition to the morphological characteristics of cellular components, three-dimentional reconstruction of nervous tissues has previously uncovered detailed cell–cell interactions such as neuron–neuron connections (e.g., synapses) and neuron–glia interactions (e.g., tripartite synapses formed by axons and astrocytic fine processes; myelin sheaths formed by axons and oligodendrocytic processes) (Tomassy et al., 2014), (Ohno et al., 2014), (Zonouzi et al., 2019). In the present study, we focused on the myelin sheaths formed by interfascicular oligodendrocytes. Of 91 myelin sheaths formed by the seven oligodendrocytes in a bead-like cluster, none shared a single axon. In other words, an axon never received multiple myelinations from a single oligodendrocyte. Although we cannot exclude the possibility that such multiple myelinations occur outside the volume we examined in the present study, the processes of an IO apparently target different axons rather than the same axon. Moreover, an oligodendrocyte appears to myelinate similar-diameter axons with a restricted range of myelin thicknesses (Fig. 3E– I). Genetic labeling of oligodendrocytes with rabies viral vector (Osanai et al., 2017), (Osanai et al., 2018) corroborated the observation that myelin fiber thicknesses (axon + myelin thickness) of IOs display statistically significant variation (Fig. 4D). The mechanisms underlying these phenomena are not clear, but recent studies may provide a clue. Oligodendrocyte processes randomly attach to and/or myelinate axons, but some of them are pruned while others form stable myelin sheaths (Czopka et al., 2013), (Almeida and Lyons, 2014). In the developmental process, IOs may form sheaths on different types of axons (in terms of diameter, activities and transmitters), but the sheaths may be selected by some unknown mechanism(s) and unstable ones removed. Alternatively, axons may secrete factors that attract certain oligodendrocytes. In any case, detailed observation of myelination processes will be required to elucidate the specific relationships between oligodendrocytic processes and axons. Recent studies suggest some mechanisms regulating neuron–oligodendrocyte coupling: in the cerebral cortex, for example, certain oligodendrocytes selectively myelinate inhibitory axons of interneurons (Zonouzi et al., 2019), implying that myelination in the CNS is not a random but a regulated event.

All reconstructed oligodendrocytes extended their processes through bundles of nearby axons and formed compact myelin around distant axons. The average linear distance was not significantly difference among the seven oligodendrocytes (Fig. 2D). Furthermore, we observed no correlation between the linear distance of oligodendrocyte processes and soma size or profile of myelination (Fig. S2). Similar results were obtained in light microscopic visualization of oligodendrocyte morphologies using RV-GFP (Fig. S3). From the viewpoint of cellular energy consumption, forming myelin sheaths on nearby axons should be more economical than extending long processes toward distant axons. On the other hand, the merit in extending long processes may be that adjacent oligodendrocytes in a beads-like cell cluster do not interfere with each other’s myelination. The mechanisms behind this apparent paradox remain enigmatic and deeper examination of the developmental processes of myelination will be required to shed light on them, as well as on the mechanisms of selective myelination discussed above. There may be common molecular and/or cellular machinery at work in these two phenomena.

We found that the myelin profiles of adjacent sheaths tended to be similar (Fig. 5). Given that a single axon rarely, if ever, receives multiple myelinations from a single oligodendrocyte (see above), adjacent sheaths will have originated from different oligodendrocytes. Myelin profiles are partly regulated by factors secreted from or expressed on the axons, exemplified by the neuregulin family (Laursen and Ffrench-Constant, 2007), (Nave and Salzer, 2006). The action potential of an axon is also an important regulating factor of myelination (Fields, 2015). Overall, adjacent myelin sheaths in the corpus callosum are likely to be under the regulation of such mechanisms.

In conclusion, we have revealed several fundamental patterns of myelination in the mouse corpus callosum, using serial three-dimensional reconstructions of electron microscopic images. The observed structural heterogeneity is likely both to contribute to the functional diversity of complex neuronal populations and to be important when modeling neuronal function.

## Acknowledgements

We thank Mrs. Atsuko Imai (National Institute for Physiological Sciences), Dr. Mariko Yamano (Nara Medical University), Dr. Yuki Terada (Nara Medical University), Dr. Ayami Isonishi (Nara Medical University), Dr. Yoshio Bando (Akita University), Mrs. Yoshie Kawabe (Nara Medical University) and Dr. Kazuki Nakahara (Nara Medical University) for technical assistance. This work was supported by a Grant-in-Aid for Scientific Research (KAKENHI) (Grant JP17H05774) from the Japan Society for the Promotion of Science, a Grant-in-Aid for Scientific Research on Innovative Areas-Resource and technical support platforms for promoting research of Advanced Bioimaging Support (JP16H06280), Research Grant from National Center of Neurology and Psychiatry (No. 30-5), Cooperative Study Programs of National Institute for Physiological Sciences, and a support from Setsuro Fujii Memorial, Osaka Foundation for Promotion of Fundamental Medical Research.

## References

Almeida RG, Lyons DA. 2014. On the resemblance of synapse formation and CNS myelination. Neuroscience 276:98–108. doi: 10.1016/j.neuroscience.2013.08.062

Baumann N, Pham-Dinh D. 2001. Biology of Oligodendrocyte and Myelin in the Mammalian Central Nervous System. Physiol Rev 81:871–927. doi: 10.1152/physrev.2001.81.2.871

Briggman KL BD. 2012. Volume electron microscopy for neuronal circuit reconstruction. Curr Opin Neurobiol 1:154–161. doi: 10.1016/j.conb.2011.10.022

Cervos-Navarro J, Artigas J, Aruffo C, Iglesias J. 1987. The fine structure of gliomatosis cerebri. Virchows Arch A Pathol Anat Histopathol 411:93–98.

Chang KJ, Redmond SA, Chan JR. 2016. Remodeling myelination: Implications for mechanisms of neural plasticity. Nat Neurosci 19:190–197. doi: 10.1038/nn.4200

Czopka T, ffrench-Constant C, Lyons DA. 2013. Individual oligodendrocytes have only a few hours in which to generate new myelin sheaths invivo. Dev Cell 25:599–609. doi: 10.1016/j.devcel.2013.05.013

Desai RA, Gao L, Raghavan S, Liu WF, Chen CS. 2009. Cell polarity triggered by cell-cell adhesion via E-cadherin. J Cell Sci 122:905–911. doi: 10.1242/jcs.028183

Dubois-Dalcq M, Behar T, Hudson L, Lazzarini RA. 1986. Emergence of three myelin proteins in oligodendrocytes cultured without neurons. J Cell Biol 102:384–392. doi: 10.1083/jcb.102.2.384

Duncan ID, Radcliff AB, Heidari M, Kidd G, August BK, Wierenga LA. 2018. The adult oligodendrocyte can participate in remyelination. Proc Natl Acad Sci U S A 201808064. doi: 10.1073/pnas.1808064115

Dupin I, Camand E, Etienne-Manneville S. 2009. Classical cadherins control nucleus and centrosome position and cell polarity. J Cell Biol 185:779–786. doi: 10.1083/jcb.200812034

Ehrengruber MU, Renggli M, Raineteau O, Hennou S, Vähä-Koskela MJV, Hinkkanen AE, Lundstrom K. 2003. Semliki Forest virus A7(74) transduces hippocampal neurons and glial cells in a temperature-dependent dual manner. J Neurovirol 9:16–28. doi: 10.1080/13550280390173346

Fields RD. 2015. A new mechanism of nervous system plasticity: Activity-dependent myelination. Nat Rev Neurosci 16:756–767. doi: 10.1038/nrn4023

Gillespie M, Stein R. 1983. The relationship between axon diameter, myelin thickness and conduction velocity during atrophy of mammalian peripheral nerves. Brain Res 259:41–56.

Haber M, Zhou L, Murai KK. 2006. Cooperative astrocyte and dendritic spine dynamics at hippocampal excitatory synapses. J Neurosci 26:8881–8891. doi: 10.1523/JNEUROSCI.1302-06.2006

Helmstaedter M, Briggman KL, Turaga SC, Jain V, Seung HS, Denk W. 2013. Connectomic reconstruction of the inner plexiform layer in the mouse retina. Nature 500:168–174. doi: 10.1038/nature12346

Hildebrand C, Hahn R. 1978. Relation between myelin sheath thickness and axon size in spinal cord white matter of some vertebrate species. J Neurol Sci 38:421–434.

Hildebrand C, Remahl S, Persson H, Bjartmar C. 1993. Myelinated nerve fibres in the CNS. Prog Neurobiol 40:319–384.

Kahn S, Tansey FA, Cammer W. 1986. Biochemical and Immunocytochemical Evidence for a Deficiency of Normal Interfascicular Oligodendroglia in the CNS of the Dysmyelinating Mutant (md) Rat. J Neurochem 47:1061–1065. doi: 10.1111/j.1471-4159.1986.tb00720.x

Kasthuri N, Hayworth KJ, Berger DR, Schalek RL, Conchello JA, Knowles-Barley S, Lee D, Vázquez-Reina A, Kaynig V, Jones TR, Roberts M, Morgan JL, Tapia JC, Seung HS, Roncal WG, Vogelstein JT, Burns R, Sussman DL, Priebe CE, Pfister H, Lichtman JW. 2015. Saturated Reconstruction of a Volume of Neocortex. Cell 162:648–661. doi: 10.1016/j.cell.2015.06.054

Laursen L, Ffrench-Constant C. 2007. Adhesion molecules in the regulation of CNS myelination. Neuron Glia Biol 3:367–375. doi: 10.1017/S1740925x08000161

Liu J, Dietz K, Deloyht JM, Pedre X, Kelkar D, Kaur J, Vialou V, Lobo MK, Dietz DM, Nestler EJ, Dupree J, Casaccia P. 2012. Impaired adult myelination in the prefrontal cortex of socially isolated mice. Nat Neurosci 15:1621–1623. doi: 10.1038/nn.3263

Maier O, Hoekstra D, Baron W. 2008. Polarity development in oligodendrocytes: Sorting and trafficking of myelin components. J Mol Neurosci 35:35–53. doi: 10.1007/s12031-007-9024-8

Marques S, Zeisel A, Codeluppi S, Van Bruggen D, Falcão AM, Xiao L, Li H, Häring M, Hochgerner H, Romanov RA, Gyllborg D, Muñoz-Manchado AB, La Manno G, Lönnerberg P, Floriddia EM, Rezayee F, Ernfors P, Arenas E, Hjerling-Leffler J, Harkany T, Richardson WD, Linnarsson S, Castelo-Branco G. 2016. Oligodendrocyte heterogeneity in the mouse juvenile and adult central nervous system. Science (80-) 352:1326–1329. doi: 10.1126/science.aaf6463

McKenzie IA, Ohayon D, Li H, De Faria JP, Emery B, Tohyama K, Richardson WD. 2014. Motor skill learning requires active central myelination. Science (80-) 346:318–322. doi: 10.1126/science.1254960

Mori S, Leblond C. 1970. Electron microscopic identification of three classes of oligodendrocytes and a preliminary study of their proliferative activity in the corpus callosum of young rats. J Comp Neurol 139:1–28.

Morizawa YM, Hirayama Y, Ohno N, Shibata S, Shigetomi E, Sui Y, Nabekura J, Sato K, Okajima F, Takebayashi H, Okano H, Koizumi S. 2017. Reactive astrocytes function as phagocytes after brain ischemia via ABCA1-mediated pathway. Nat Commun 22:28. doi: 10.1038/s41467-017-00037-1

Nave K, Salzer J. 2006. Axonal regulation of myelination by neuregulin 1. Curr Opin Neurobiol 16:492–500.

Nimmerjahn A, Kirchhoff F, Helmchen F. 2005. Neuroscience: Resting microglial cells are highly dynamic surveillants of brain parenchyma in vivo. Science (80-) 308:1314–1318. doi: 10.1126/science.1110647

Nimmerjahn A, Kirchhoff F, Kerr J, Helmchen F. 2004. Sulforhodamine 101 as a specific marker of astroglia in the neocortex in vivo. Nat Methods 1:31–37.

Ogawa T, Hagihara K, Suzuki M, Yamaguchi Y. 2001. Brevican in the developing hippocampal fimbria: differential expression in myelinating oligodendrocytes and adult astrocytes suggests a dual role for brevican in central nervous system fiber tract development. J Comp Neurol 432:285–295.

Ohno N, Chiang H, Mahad DJ, Kidd GJ, Liu L, Ransohoff RM. 2014. Mitochondrial immobilization mediated by syntaphilin facilitates survival of demyelinated axons. Proc Natl Acad Sci U S A 111:9953–9958. doi: 10.1073/pnas.1401155111

Ohno N, Kidd GJ, Mahad D, Kiryu-Seo S, Avishai A, Komuro H, Trapp BD. 2011. Myelination and Axonal Electrical Activity Modulate the Distribution and Motility of Mitochondria at CNS Nodes of Ranvier. J Neurosci 31:7249–7258. doi: 10.1523/JNEUROSCI.0095-11.2011

Osanai Y, Shimizu T, Mori T, Hatanaka N, Kimori Y, Kobayashi K, Koyama S, Yoshimura Y, Nambu A, Ikenaka K. 2018. Length of myelin internodes of individual oligodendrocytes is controlled by microenvironment influenced by normal and input-deprived axonal activities in sensory deprived mouse models. Glia 66:2514–2525. doi: 10.1002/glia.23502

Osanai Y, Shimizu T, Mori T, Yoshimura Y, Hatanaka N, Nambu A, Kimori Y, Koyama S, Kobayashi K, Ikenaka K. 2017. Rabies Virus-Mediated Oligodendrocyte Labeling Reveals a Single Oligodendrocyte Myelinates Axons from Distinct Brain Regions. Glia 65:93–105. doi: 10.1002/glia.23076

Peters A, Palay S, Webster H. 1991. The fine structure of the nervous system. Neurons and their supporting cells. Oxford University Press.

Sampaio-Baptista C, Johansen-Berg H. 2017. White Matter Plasticity in the Adult Brain. Neuron 96:1239–1251. doi: 10.1016/j.neuron.2017.11.026

Simons M, Nave KA. 2016. Oligodendrocytes: Myelination and axonal support. Cold Spring Harb Perspect Biol 8:1–16. doi: 10.1101/cshperspect.a020479

Snaidero N, Möbius W, Czopka T, Hekking LHP, Mathisen C, Verkleij D, Goebbels S, Edgar J, Merkler D, Lyons DA, Nave KA, Simons M. 2014. Myelin membrane wrapping of CNS axons by PI(3,4,5)P3-dependent polarized growth at the inner tongue. Cell 156:277–290. doi: 10.1016/j.cell.2013.11.044

Suzuki M, Raisman G. 1992. The glial framework of central white matter tracts: segmented rows of contiguous interfascicular oligodendrocytes and solitary astrocytes give rise to a continuous meshwork of transverse and longitudinal processes in the adult rat fimbria. Glia 6:222–235.

Tomassy GS, Berger D, Chen H-H, Kasthuri N, Hayworth K, Vercelli A, Seung S, Lichtman J, Arlotta P. 2014. Distinct Profiles of Myelin Distribution. Science (80-) 344:319–324. doi: 10.1126/science.1249766

Wake H, Lee PR, Fields RD. 2011. Control of local protein synthesis and initial events in myelination by action potentials. Science (80-) 333:1647–1652. doi: 10.1126/science.1206998

Yamasaki R, Lu H, Butovsky O, Ohno N, Rietsch AM, Cialic R, Wu PM, Doykan CE, Lin J, Cotleur AC, Kidd G, Zorlu MM, Sun N, Hu W, Liu L, Lee J, Taylor SE, Uehlein L, Dixon D, Gu J, Floruta CM, Zhu M, Charo IF, Weiner HL, Ransohoff RM. 2014. Differential roles of microglia and monocytes in the inflamed central nervous system. J Exp Med 211:1533–1549. doi: 10.1084/jem.20132477

Yamazaki Y, Fujiwara H, Kaneko K, Hozumi Y, Xu M, Ikenaka K, Fujii S, Tanaka KF. 2014. Short- and long-term functional plasticity of white matter induced by oligodendrocyte depolarization in the hippocampus. Glia 62:1299–1312. doi: 10.1002/glia.22681

Zonouzi M, Berger D, Jokhi V, Kedaigle A, Lichtman J, Arlotta P, Zonouzi M, Berger D, Jokhi V, Kedaigle A, Lichtman J, Arlotta P. 2019. Individual Oligodendrocytes Show Bias for Inhibitory Axons in the Neocortex. Cell Rep 27:2799-2808.e3. doi: 10.1016/j.celrep.2019.05.018

